# Behavioral characteristics of an extremely old rhesus macaque in a zoo: Dementia-like symptoms and implications for quality of life of geriatric animals

**DOI:** 10.64898/2026.03.17.712497

**Authors:** Yumi Yamanashi, Haruna Bando, Keita Niimi, Daisuke Nakagawa, Susumu Iwaide, Tomoaki Murakami

## Abstract

Documenting and understanding the welfare of aging animals are crucial for maintaining their well-being and making appropriate management decisions. This study details the behaviors of an extremely old rhesus macaque (ISK) in which senile plaques and phosphorylated tau deposition were observed in post-mortem pathological analyses of the brain. We report on the activity bsudgets, behavioral rhythms, gait, quality of life (QoL) scores, and anecdotal episodes of this individual. The average 24-hour activity budgets, analyzed from surveillance camera recordings, revealed that ISK spent most of her time inactive. ISK was sometimes active at night, though her behavior remained predominantly diurnal. Gait analysis suggested that her movement patterns changed between the first (December 2020) and the last (June 2021) assessment. QoL assessments, using a scoring sheet, indicated relatively good well-being until the later stage of her life. An anecdotal episode, along with the husbandry diary, suggested signs of cognitive decline. These results suggest possible signs of physical decline, and some behavioral changes that could be associated with cognitive decline in an extremely old rhesus macaque. However, we could not confirm cognitive dysfunction without further controlled cognitive testing. We hope that future studies will consider the behavioral symptoms observed in this study as monitoring items to better understand physical and cognitive decline, and possible relationships with QoL in primates.

## Introduction

Documenting and understanding the welfare of aging animals are crucial for maintaining their well-being and making appropriate management decisions (Brando & Chapman, 2023). It also leads to an understanding of the species-specific aging process. Primates follow distinct life cycles that are characterized by a long developmental period and longevity. A relatively large brain and high cognitive abilities are also typical (Primate Society of Japan, 2023). Studies on primate aging have revealed the cognitive and physical changes associated with aging (e.g. Baxter, Roberts, Roberts, & Rapp, 2023; Bliss-Moreau & Baxter, 2019; Chaudron, Pifferi, & Aujard, 2021; Lacreuse et al., 2005; Lacreuse, Raz, Schmidtke, Hopkins, & Herndon, 2020; Lacreuse, Russell, Hopkins, & Herndon, 2014; Languille et al., 2012; Moore, Killiany, Herndon, Rosene, & Moss, 2006; Némoz-Bertholet & Aujard, 2003). Several pathological reports have suggested the occurrence of Alzheimer’s disease-like lesions in aged primates (Nakamura et al., 1998; Paspalas et al., 2018; Uno & Walker, 1993), although the pathological processes may not be completely identical between humans and non-human primates. For example, the post-mortem pathological analysis of a 43-year-old rhesus macaque revealed similarities to human patients with Alzheimer’s disease, including the aggregation of amyloid β (Aβ) in the form of senile plaques and phosphorylated tau proteins (Iwaide et al., 2023). However, pathological cognitive decline in nonhuman primates has rarely been observed (Finch & Austad, 2012). A recent study suggested an association between tau pathology and neurological symptoms (parkinsonism) in a cynomolgus macaque; however, cognitive decline was not documented (Takahashi et al., 2026). A study on gray mouse lemurs suggested declines in object discrimination learning (Hozer, Pifferi, Aujard, & Perret, 2019). It also suggested that the most extreme outlier in cognitive performance was the subject with the highest cortical and extracellular Aβ burden, including in the hippocampus. The authors further discussed that these outliers may represent cases of pathological brain aging.

Dementia is well studied in humans, dogs, and cats (Chambers et al., 2015; Chapagain, Range, Huber, & Virányi, 2017; Gunn-Moore, 2011; Madari et al., 2015; Ozawa, 2020; Szabó, Gee, & Miklósi, 2016). In humans, symptoms include the loss of abilities in several cognitive domains (language, visuospatial, executive, or other abilities), the impairment of functional abilities for day-to-day life (e.g., social, occupational, and self-care), psychological changes, altered sleep/wake cycle, and gait impairment (Arvanitakis, Shah, & Bennett, 2019), all of which can affect quality of life (QoL) (Krivanek, Gale, McFeeley, Nicastri, & Daffner, 2021). In canine and feline species, behavioral changes such as spatial disorientation or confusion (D: disorientation), altered social relationships (I: interaction), altered behavioral responses to stimuli, changes in sleep/wake patterns (S: sleep–wake cycle), house-soiling (H: house-soiling), altered learning and memory, changes in activity (A: activity), altered interest in food, and temporal disorientation have been observed (Gunn-Moore, 2011; Madari et al., 2015; Salvin, McGreevy, Sachdev, & Valenzuela, 2011). The acronym DISHA is used to highlight the different categories of symptoms (Benzal & Rodríguez, 2016). Such symptoms, which can be observed through daily interactions with animals, are important when considering their daily care. However, comparable reports of behavioral symptoms in nonhuman primates are currently lacking (Walker & Jucker, 2017). Furthermore, studies integrating individual behavioral observations with post-mortem neuropathological findings remain limited.

This study details the behaviors of the individual reported in Iwaide et al. (2023). This individual was unique owing to the old age and the extent of hyperphosphorylated tau deposition and small amounts of neurofibrillary tangles observed in the hippocampus. Paspalas et al. (2018) reported tau aggregation but no behavioral symptoms, except for impaired performance on a delayed nonmatch-to-sample task of recognition memory in aged rhesus macaques. We report the activity budgets, behavioral rhythms, gait, quality of life (QoL) scores, and anecdotal episodes of this individual, and compare some parameters with those of other individuals housed in the same location, to identify potential behavioral associations with the pathological changes.

## Methods

The main subject was a female rhesus macaque who lived up to 43 years and 4 months (hereafter referred to as ISK) and died without euthanasia on September 12, 2021 at the Kyoto City Zoo, Japan (Supplementary table 1). In 2021, she was registered as the longest-living rhesus macaque by Guinness World Records (https://www.guinnessworldrecords.com/), older than rhesus macaques in previous studies (e.g. 40 years old: Finch & Austad, 2012). From age 41, the keepers noticed behavioral changes and recorded noteworthy observations in the husbandry diary on an ad hoc basis. These changes, along with potential similarities to symptoms of cognitive decline in dogs and cats, are summarized in Supplementary table 2. ISK used both outdoor (approximately 200 m^2^) and indoor (approximately 37 m^2^) enclosures until January 2021, but primarily used the indoor enclosure beyond that date after sustaining frequent injuries in the outdoor enclosure. Keepers occasionally took her to the outdoor enclosure under supervision, even after January 2021, to allow her to explore the environment with her conspecifics. Both the outdoor and indoor enclosures contained 3D structures for the monkeys to climb. The outdoor enclosures also contained various types of vegetation for exploration and foraging. Feeding occurred at least three times per day. From approximately 9:00 am–5:00 pm, the lights were on, and natural sunlight also entered the facility. The keepers switched off the lights at approximately 5:00 pm each day. ISK lived with two female conspecifics until 2 days before her death.

We also described the gait and nocturnal activity rhythms of 14 monkeys (Supplementary table1: aged 14 to 37 years). Two of the 14 monkeys (32- and 37-year-old females) lived together with ISK (geriatric group monkeys). The remaining 12 monkeys (aged 14 to 26 as of May 2023) began using the same indoor and outdoor enclosures in November 2022, after the geriatric group monkeys left the enclosure.

A surveillance camera (AXIS 1065-LW) was installed in the rhesus macaque indoor enclosure. We recorded ISK’s general behaviors for a total of 551 hours between January and August 2021, based on 24-hour video recordings collected on 23 days (2–3 days per month; one recording hour was unavailable because of technical issues). Behavioral data were extracted from the full 24-hour recordings at 5-minute intervals (Yoshida, Tanaka, & Wada, 2017). Sample days were selected based on the availability of usable video recordings. We recorded whether the animal was engaging in inactive, moving, eating, exploring, social behaviors (agonistic interaction/ grooming), or other behaviors. We recorded a timeout if we could not see her due to the camera angle. To record night-time activity rhythms of 12 other monkeys (between 14 and 26 years old), we collected overnight video recordings between April and August 2023 for a total of 108 hours (18:00-06:00 across 9 nights). We scored all recorded overnight videos using a scan sampling method and recorded the number of animals exhibiting active behavior (Martin & Bateson, 2007). Active behaviors included moving, eating, exploring, or social behaviors (agonistic interaction/grooming). YY, the first author, collected all behavioral data.

Gait analyses were performed on the video data collected between December 2020 and June 2021 (9 video clips) for ISK and between 2020 and 2023 for other 14 individuals (25 video clips). We only used videos with clear and unobstructed views of limb touchdowns (Bernstein-Kurtycz et al., 2023). To analyze ISK’s gait in various contexts, we used pre-recorded videos that, although not primarily intended for gait analysis, provided clear views of the gait action. We calculated relative stride length, mean stride duration, relative speed, limb phase (i.e., the proportion of stride duration separating hindlimb touchdown from ipsilateral forelimb touchdown during symmetrical gaits), the mean number of supporting limbs, and duty factor (i.e., the amount of time a limb was in contact with the ground divided by stride duration) following the previous study(Bernstein-Kurtycz et al., 2023). We also calculated the ipsilateral duty factor from duty factors in accordance with Bernstein-Kurtycz et al. (2023), namely ((L_df_ − R_df_) / (0.5 × (L_df_ + R_df_)) × 100, where L_df_ is the mean of left forelimb and left hindlimb duty factors and R_df_ is the mean of right forelimb and right hindlimb duty factors. Values close to 0 indicate little or no left-right asymmetry, whereas positive values indicate relatively greater duty factors on the left side and negative values indicate relatively greater duty factors on the right side; larger absolute values indicate greater asymmetry. We predicted that the individual would exhibit slower speed, shorter stride length, and altered gait patterns, including impaired coordination, as reflected in parameters such as limb phase, the mean number of supporting limbs, duty factor, and ipsilateral duty factor, compared with the other individuals. Although previous studies recorded videos mostly using the same cameras (Bernstein-Kurtycz et al., 2023; Dunham, McNamara, Shapiro, Hieronymus, & Young, 2018; Dunham et al., 2019; Dunham, McNamara, Shapiro, Phelps, & Young, 2020), we used videos recorded from different cameras (the list of cameras is presented in the Supplementary table 3). To check the consistency of video recordings, we recorded gaits of 6 occasions using two cameras (GoPro HERO9 Black and Hykecam LT4G) and analyzed the gaits using GaitKeeper2 (Dunham et al., 2018). We obtained consistent results from the recordings collected with different cameras, as indicated by a small standard deviation within the same gait event and a larger standard deviation between different gait events (Supplementary table 3).

QoL analyses were conducted using a modified QoL assessment sheet developed by Mandai Wildlife Group. A veterinarian (DN) monitored and recorded animals’ QoL from January to September 2021. The assessment sheet consisted of six areas: pain, maintenance behaviors, appetite, locomotion, object manipulation/exploration, and interaction with the environment; each was rated on a five-point scale, with a score one indicating a severe condition and a score of five indicating a good condition (Supplementary table 4). At Kyoto City Zoo, QoL scoring was conducted primarily for selected animals with impaired physical or mental health, at intervals deemed appropriate by zoo staff. This study was carried out during the early phase of implementation of this QoL assessment system.

Because some unusual behaviors were observed in ISK, a detailed episode of the unusual behaviors that were observed by YY on December 28, 2020 is also provided.

### Statistical analyses

R 4.4.1 was used for statistical analyses and data visualization (R Development Core Team, 2023). We divided the time range into two sections: diurnal (6:00 am– 5:55pm) and nocturnal (6:00 pm–5:55 am the next morning). We compared ISK’s rate of active behaviors between diurnal and nocturnal time periods using the Wilcoxon signed-rank test and the wilcox.exact function from the ‘exactRankTests’ package (Hothorn, Hornik, van de Wiel, & Zeileis, 2006). To compare the differences in nocturnal behavioral rhythms between ISK and other individuals, we visualized the data to determine whether ISK’s behaviors fall within the range of those of other individuals. We visualized the data using ggplot2 (Wickham, 2016). We calculated the rate of active behaviors in each 1-hour time period (e.g., 18:00–18:55) and used it for subsequent analyses. We calculated z-scores as (*Mi − Mo*) / *SDo*, where Mi is the mean of ISK’s data, Mo is the mean of the other individuals’ data, and SDo is the standard deviation of the other individuals’ data. We considered ISK’s data to be different from those of the other individuals if ∣z∣ was greater than 2 (DeVore, 2017). Gait differences between ISK and the other individuals were analyzed using Z-scores, as described above. The data that support the findings of this study are openly available in Open Science Framework (Yamanashi, 2024).

## Results

### Activity budgets

The average 24-hour activity budgets across the periods revealed that ISK spent most of the time inactive (Average ± SD: 89.1 ± 3.06 % between January and April, 94.9 ± 2.65 % between May and August) but was occasionally observed eating (Average ± SD: 5.58 ± 1.73 % between January and April, 3.80 ± 1.56 % between May and August), exploring (Average ± SD: 1.78 ± 1.55 % between January and April, 0.12 ± 0.37 % between May and August), moving (Average ± SD: 3.33 ± 2.22 % between January and April, 1.10 ± 1.26 % between May and August), and engaging in social behaviors (Average ± SD: 0.084 ± 0.29 % between January and April, 0.00 ± 0.00 % between May and August).

The activity budgets changed from January to August: the rate of inactivity increased and the active behaviors (e.g., exploring and moving) decreased (Figure 1). In January, we observed ISK grooming another individual for a short period of time, but this behavior was not subsequently observed. We also noted that ISK reacted to the behaviors of others (e.g., attempting to protect food from other animals) in the latter part of the study, although we were unable to quantify this using the current methodology (time-sampling).

**Figure 1.**
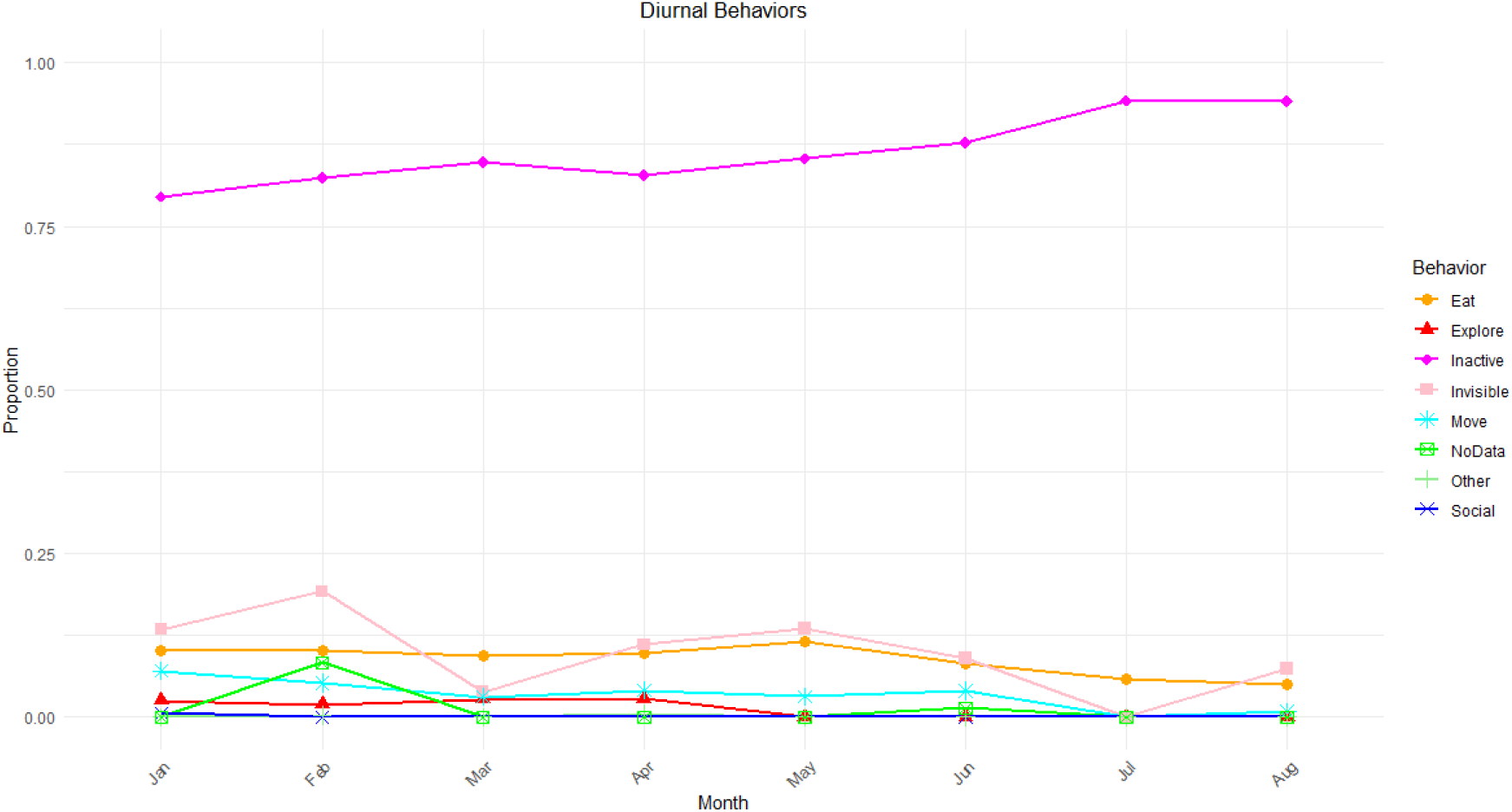
Changes in the diurnal activity budgets in the final 8 months.

### Behavioral rhythm

ISK’s behavior remained primarily diurnal (Figure 2: V = 276, p < 0.001), although she tended to be more active than the other monkeys at night (Figure 3). We could not calculate the z-score for the time period (T22: 22:00–22:55) because all data points for the other individuals were 0.

**Figure 2.**
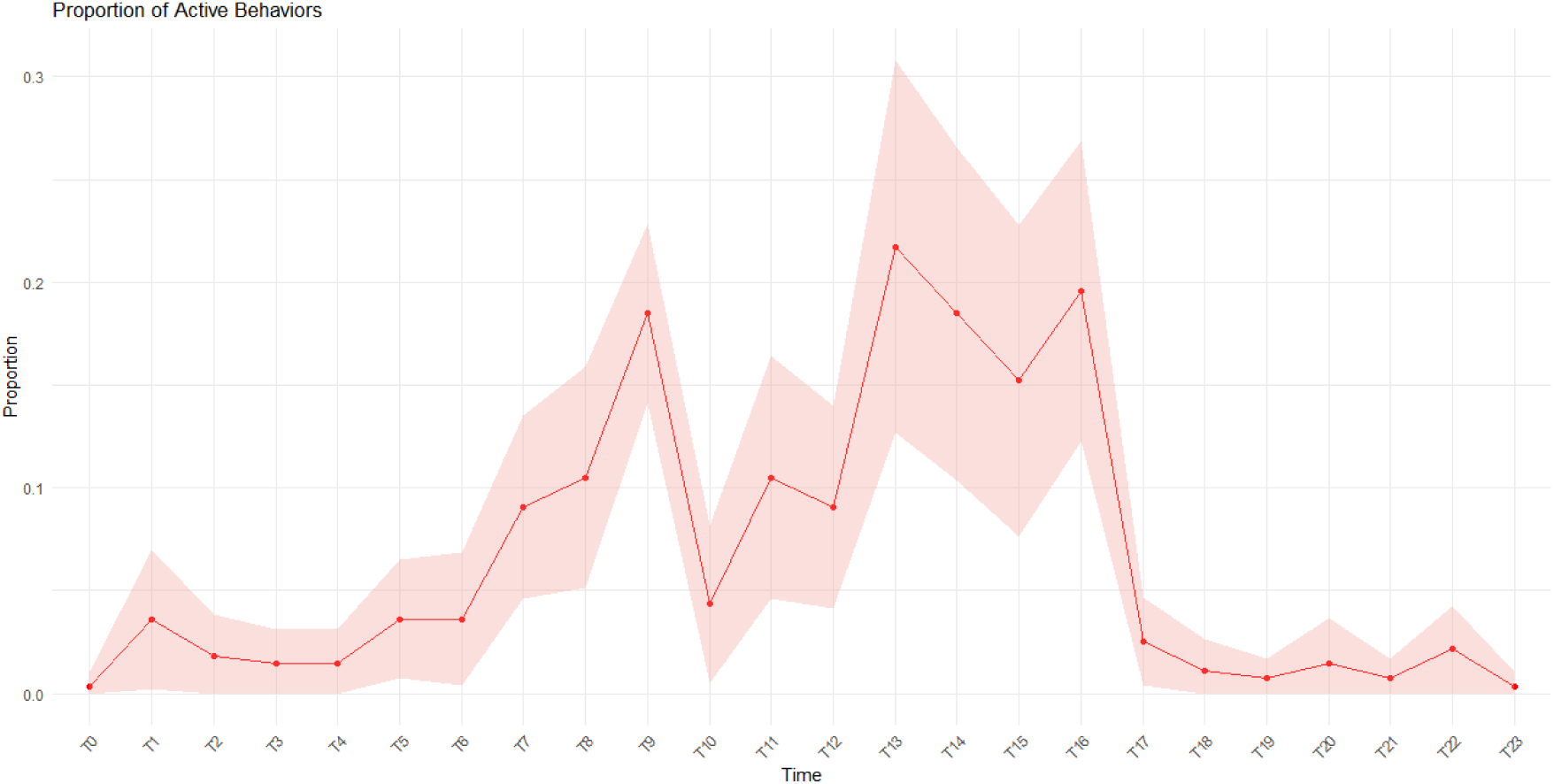
24hrs activity patterns with 95% confidence intervals. The red line represents the mean proportion of active behaviors at each time point, while the shaded red region represents the 95% confidence interval (CI).

**Figure 3.**
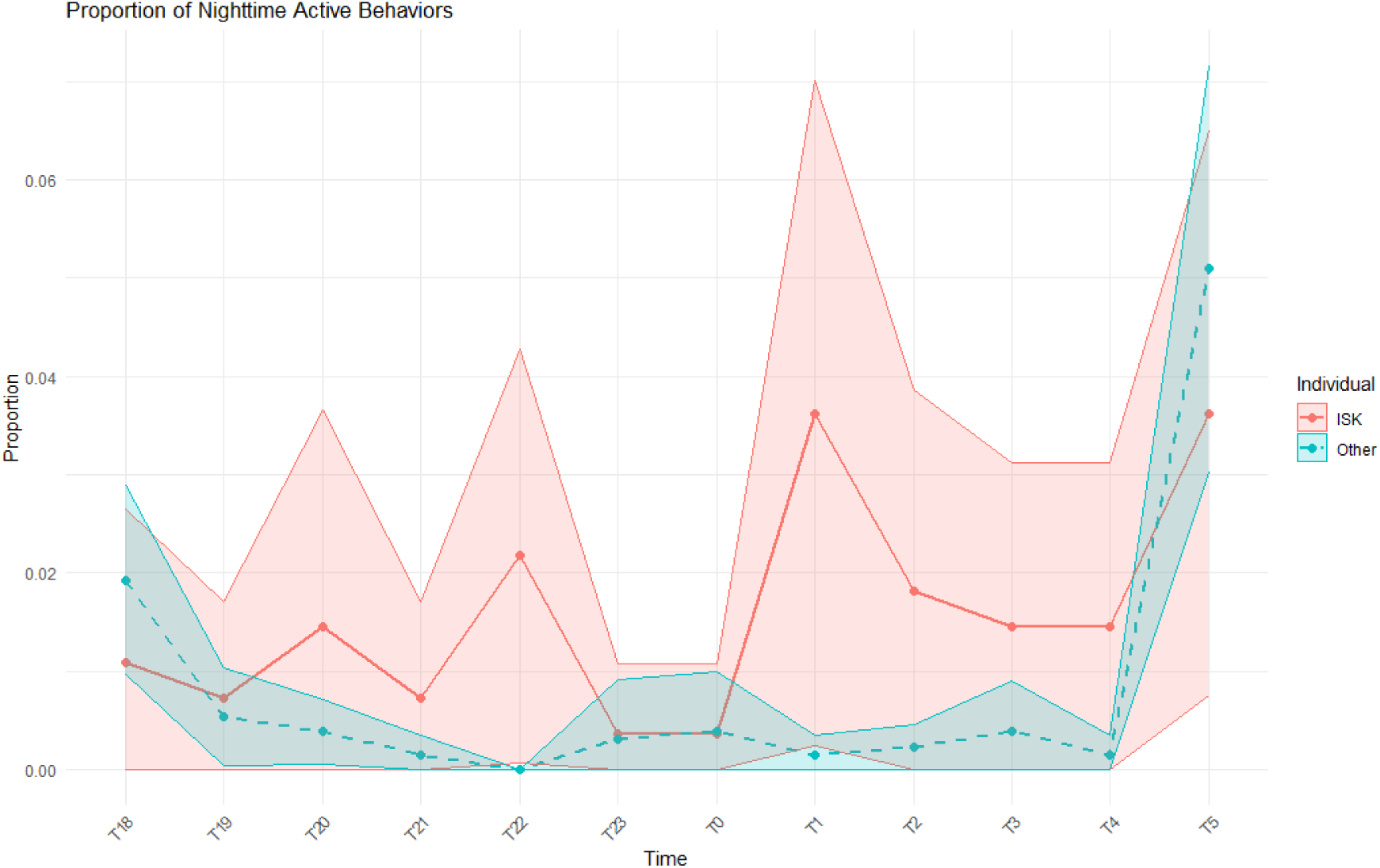
Nighttime behaviors of ISK and other individuals The red line represents ISK’s mean proportion of active behaviors over time, with the shaded red region indicating the 95% confidence interval (CI). The blue dashed line represents the mean proportion of active behaviors for other individuals, with the shaded blue region indicating the 95% CI. The asterisks indicate the points where the z-score was greater than 2.

### Gait analyses

Gait analysis revealed that ISK exhibited variations in multiple parameters when comparing the earliest (December 2020) and latest (June 2021) data. Additionally, ISK exhibited slower movements, a greater ipsilateral duty factor, and a greater left limb phase at touchdown and right limb phase at midstance compared to the other individuals (Table 1).

**Table 1.**
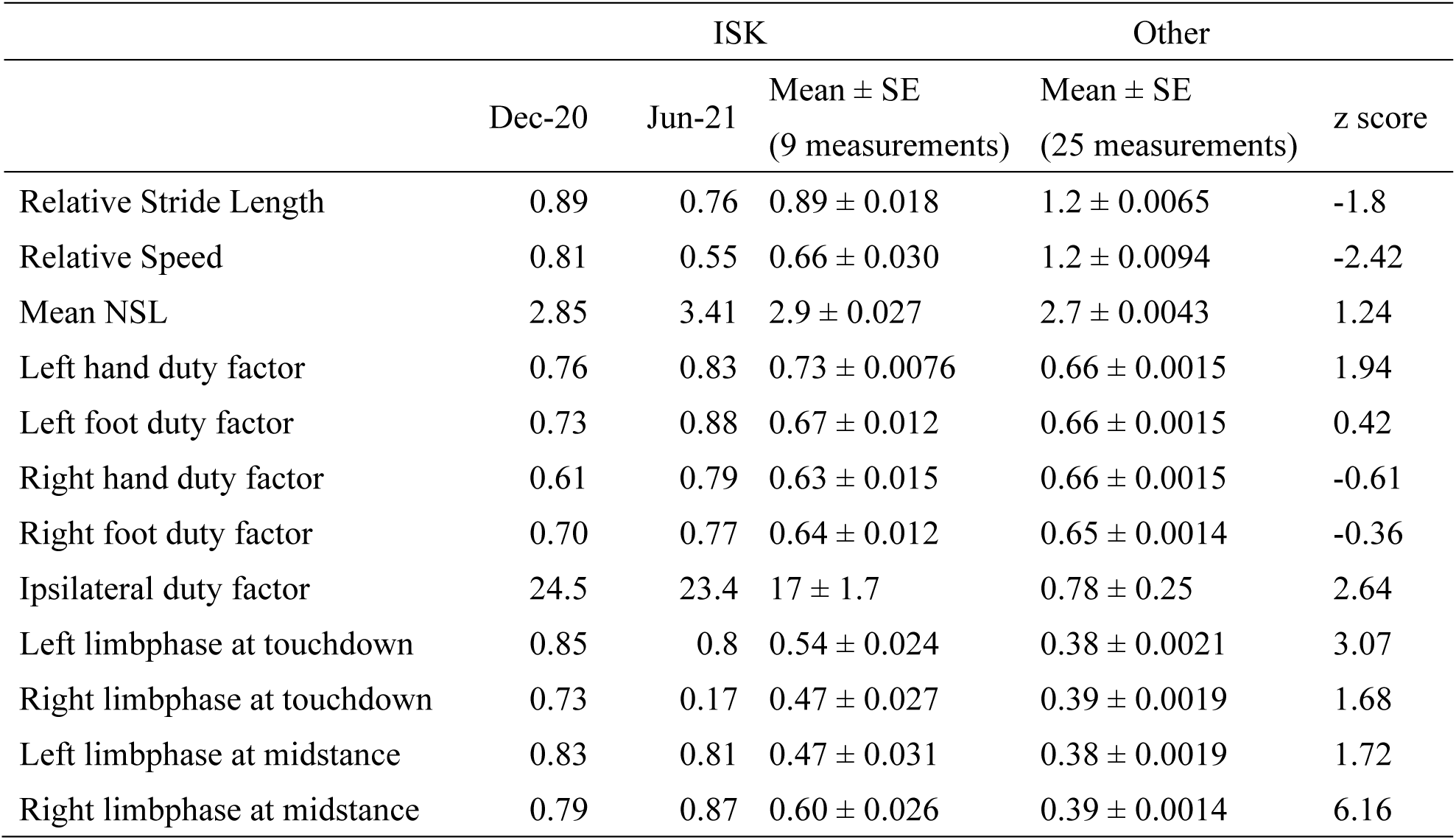
Gait characteristics of ISK.

|  | ISK |  |  | Other | z score |
| --- | --- | --- | --- | --- | --- |
| | Dec-20 | Jun-21 | Mean ± SE<br>(9 measurements) | Mean ± SE<br>(25 measurements) | |
| Relative Stride Length | 0.89 | 0.76 | $0.89 \pm 0.018$ | $1.2 \pm 0.0065$ | -1.8 |
| Relative Speed | 0.81 | 0.55 | $0.66 \pm 0.030$ | $1.2 \pm 0.0094$ | -2.42 |
| Mean NSL | 2.85 | 3.41 | $2.9 \pm 0.027$ | $2.7 \pm 0.0043$ | 1.24 |
| Left hand duty factor | 0.76 | 0.83 | $0.73 \pm 0.0076$ | $0.66 \pm 0.0015$ | 1.94 |
| Left foot duty factor | 0.73 | 0.88 | $0.67 \pm 0.012$ | $0.66 \pm 0.0015$ | 0.42 |
| Right hand duty factor | 0.61 | 0.79 | $0.63 \pm 0.015$ | $0.66 \pm 0.0015$ | -0.61 |
| Right foot duty factor | 0.70 | 0.77 | $0.64 \pm 0.012$ | $0.65 \pm 0.0014$ | -0.36 |
| Ipsilateral duty factor | 24.5 | 23.4 | $17 \pm 1.7$ | $0.78 \pm 0.25$ | 2.64 |
| Left limbphase at touchdown | 0.85 | 0.8 | $0.54 \pm 0.024$ | $0.38 \pm 0.0021$ | 3.07 |
| Right limbphase at touchdown | 0.73 | 0.17 | $0.47 \pm 0.027$ | $0.39 \pm 0.0019$ | 1.68 |
| Left limbphase at midstance | 0.83 | 0.81 | $0.47 \pm 0.031$ | $0.38 \pm 0.0019$ | 1.72 |
| Right limbphase at midstance | 0.79 | 0.87 | $0.60 \pm 0.026$ | $0.39 \pm 0.0014$ | 6.16 |

### A potential episode of cognitive decline

On December 28, 2020, 09:36, video recording started. ISK was resting on the ladder located around area A with her two conspecifics (Figure 4). 09:55, ISK moved and foraged area A. 09:56, ISK explored at area A. 10:01, ISK walked around areas B–C. 10:03, ISK walked to the ground area (area D). 10:06, ISK explored and ate grasses and soils (area D). This continued for more than 13 mins. 10:19, ISK walked to the bush and became entangled in vines, but still moved forward. 10:22, ISK continued to move forward entangled in vines, but sometime rested with her eyes closed and approached the dry moat. 10:23, ISK continued to move forward and fell into the dry moat, which was approximately 50 cm high and 48 cm wide (area E). 10:25, ISK was rescued by a keeper. The movie was included in the supplementary materials.

**Figure 4.**
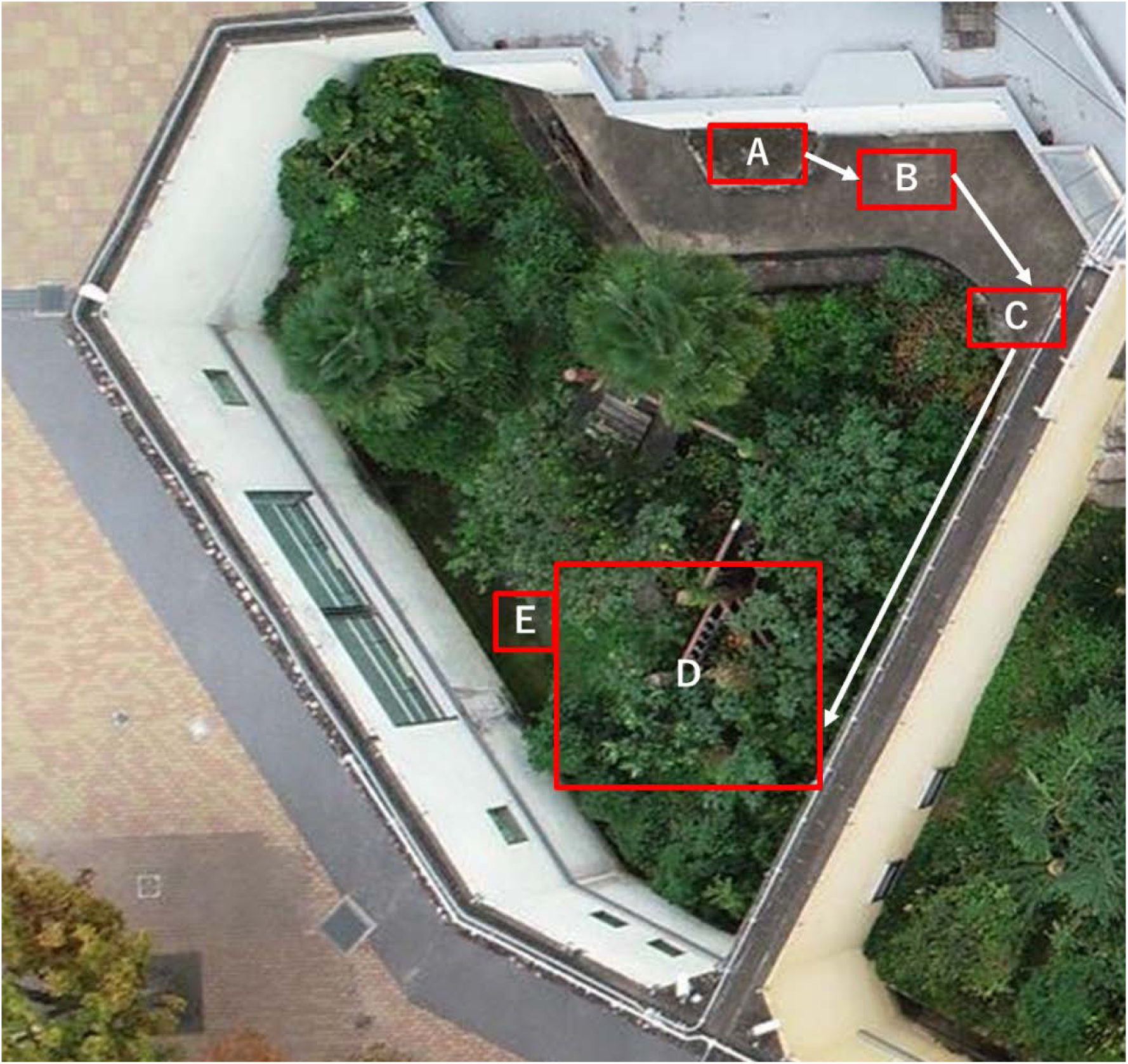
The locations inside the outdoor enclosure.

### QoL assessment

QoL was relatively well maintained until the later stages of her life (Figure 5). The results of behavioral characteristics are summarized in Table 2.

**Figure 5.**
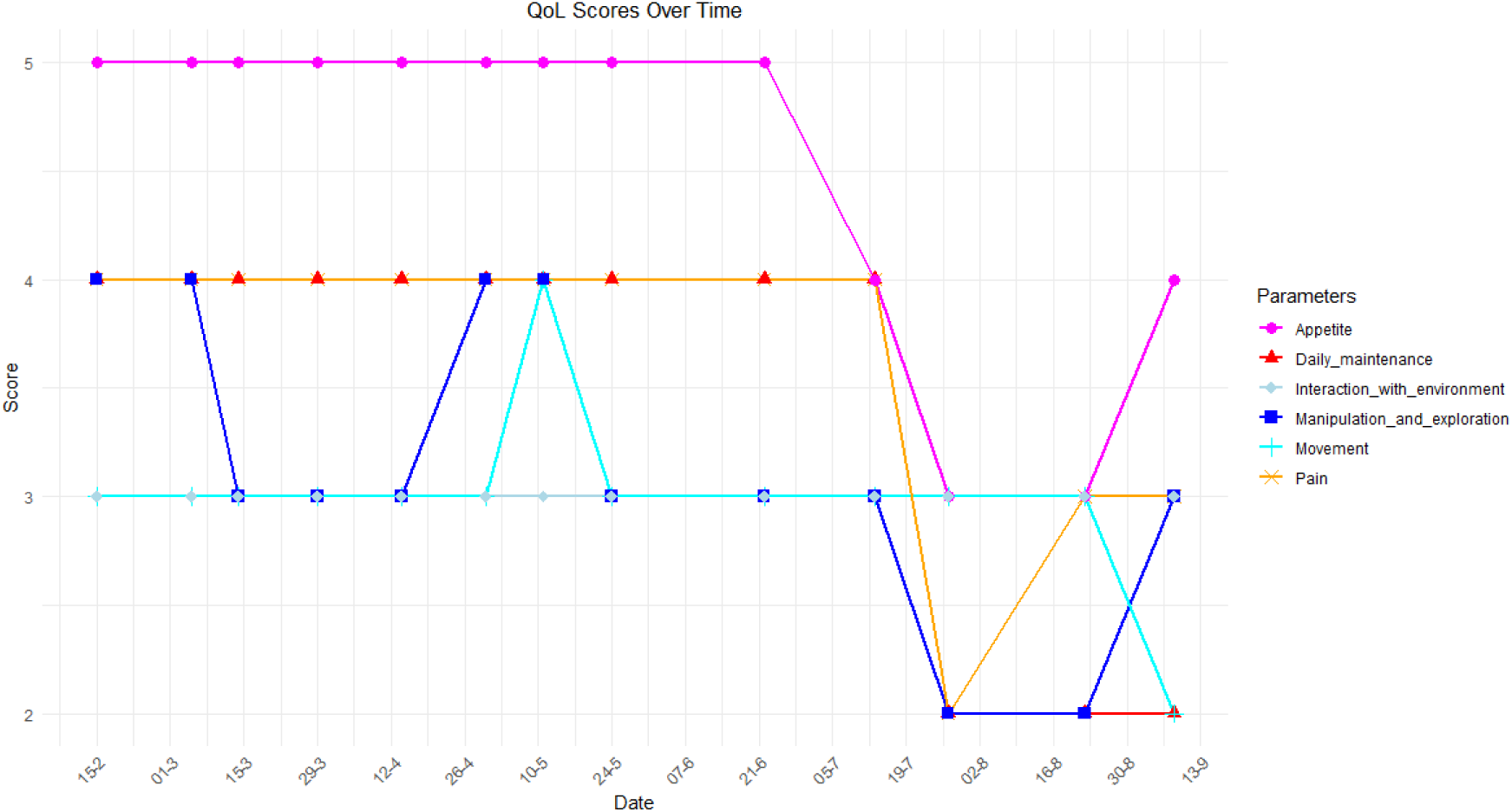
Changes in QoL scores in the final 8 months.

**Table 2.**
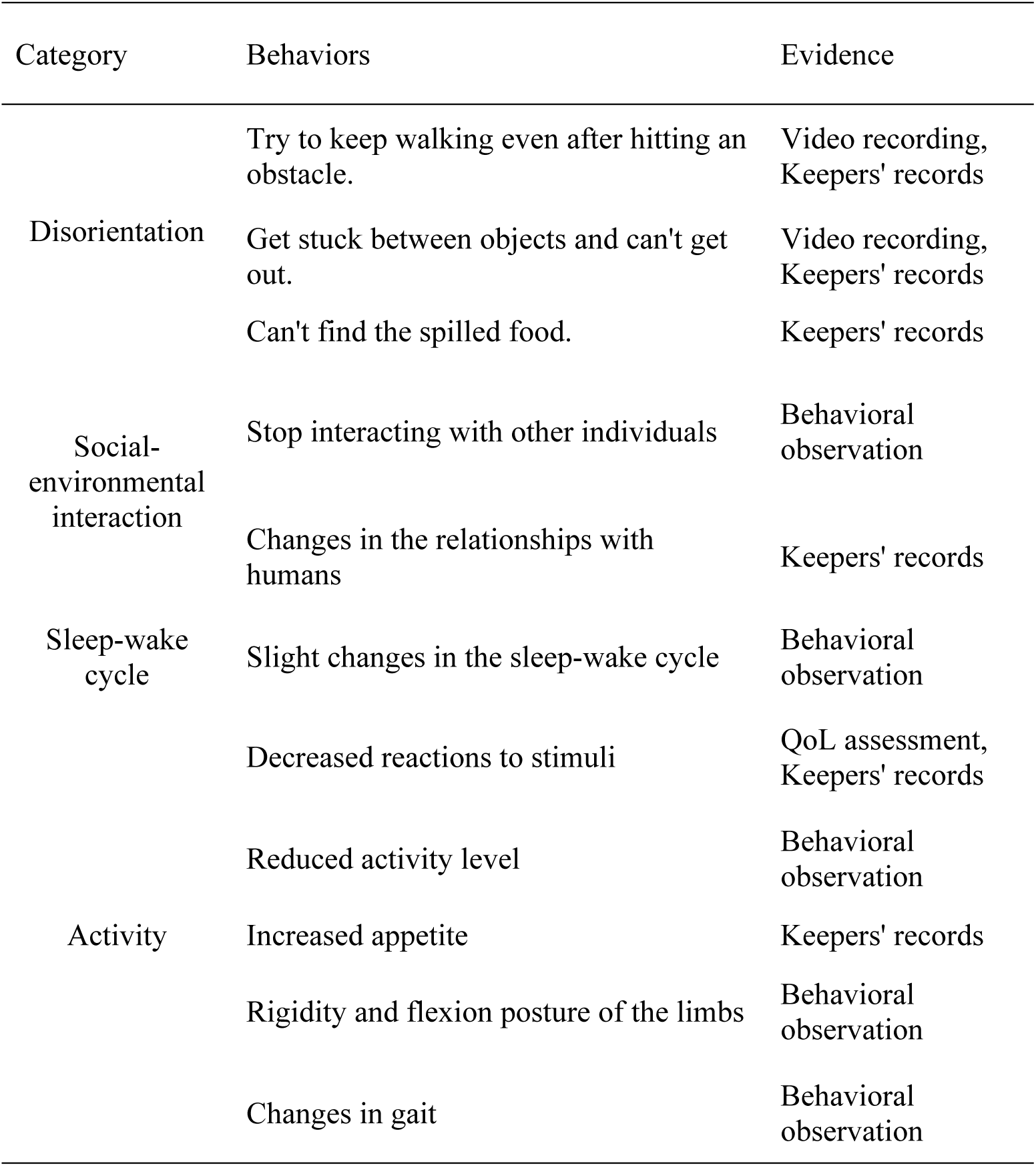
Summary of behavioral characteristics.

## Discussion

We have presented a rare documentation of the behavioral characteristics of an extremely old rhesus macaque. We observed some similarities to canine and feline cognitive dysfunction (Table 2: e.g., getting stuck between objects and being unable to escape, or continuing to walk even after hitting an obstacle.). Post-mortem pathological analyses revealed severe Aβ deposition, hyper phosphorylated tau deposition and small amounts of neurofibrillary tangles in the hippocampus of ISK (Iwaide et al., 2023). The same analyses on other elderly individuals who died at the ages of 37 and 32 at the Kyoto City Zoo did not find hyperphosphorylated tau deposition or signs of cognitive decline. Taken together, these physiological and behavioral observations suggest that ISK may have been experiencing age-related cognitive changes, potentially involving Alzheimer’s disease-like processes. The pathological processes may not be completely identical between humans and non-human primates. For example, tau aggregation is considered to be a cause of cognitive dysfunction in humans (Takeda, 2019), but only small amounts of neurofibrillary tangles were observed in ISK, despite her advanced age. Furthermore, we were not able to directly link tau-related pathology with cognitive decline through cognitive testing as in some previous studies (Baxter et al., 2023; Chaudron et al., 2021; Schmidtke et al., 2020). In addition, ageing likely progresses differently among individuals, and not all expected signs of cognitive decline were observed in ISK. For example, mild changes in the sleep–wake cycle were observed in ISK (higher nocturnal activity than other individuals), but she primarily maintained diurnal patterns. The changes in gait could have been related to both physical and cognitive decline. Behaviors suggestive of disorientation were also recorded mainly in the complex outdoor enclosure and not in the indoor enclosure, indicating that environmental context may have influenced their expression. Nevertheless, the strength of this case study lies in the combination of physiological findings and longitudinal behavioral observations, which are rarely available together in studies of ageing in non-human primates. Although our observations alone are insufficient to conclusively document cognitive decline, the overall pattern of changes in this exceptionally old individual suggests that such a process may have been occurring. Future studies combining longitudinal behavioral observations with systematic cognitive testing in ageing individuals will be necessary to clarify whether these pathological and behavioral changes are reliably associated with cognitive decline in non-human primates. However, it is important to note that these diary records began when ISK was 41 years old, which is far beyond the average life expectancy of rhesus macaques (Arnsten, Datta, & Preuss, 2021). Documenting such rare cases in primates will be essential for improving our understanding of age-related cognitive changes and their pathological correlates.

QoL scores remained relatively high just before ISK’s death, although we observed various symptoms of physical and cognitive dysfunction. She remained in a social group (only with two old individuals who had a good relationship with her) until two days before her death. The rate of interactions with other conspecifics was very low, but she did not experience any aggressive behavior from them and occasionally vocalized in response to their vocalizations. She did not show any atypical behaviors nor depressive-like responses that were observed in a previous study of a geriatric primate (Simpson, Grove, Bronson, & Herrelko, 2024). Combined with the limited observable behavioral changes, the pathological changes in her brain, such as Aβ burden and hyperphosphorylated tau deposition, did not severely deteriorate her QoL. The brain aging process of non-human primates is milder compared to that in humans and may not have a significant impact (Isidro, 2024). This may also be due to the keepers and veterinarians supporting her life by modifying her diet, environment, and care based on her symptoms.

The current study utilized multiple approaches to objectively monitor ISK’s condition. For the quantitative behavioral analyses, we used a surveillance camera system to analyze her behaviors in detail and identify subtle changes (Yoshida et al., 2012). As quantitative behavioral recording is a time-consuming methodology, it is also important to have simple and easy assessment tools that can be used on-site. Comparisons of diurnal behavioral data with QoL scores revealed their similarity: QoL scores decreased in July when the rate of inactivity increased and the rates of movement and eating decreased. Nocturnal behaviors were already dominated by inactivity and did not correspond to the changes in QoL scores. Gait analysis suggested that her movement patterns changed between the first (December 2020) and the last (June 2021) assessment. Comparative gait analysis revealed that ISK’s gait was different from that of other individuals. Not only did her moving speed decrease, as also suggested in the previous study (Shively et al., 2012), but other parameters, such as limb phase and duty factors, also differed from those of normal individuals with a stable gait. This suggests that her locomotion ability and gait coordination were impaired. QoL scores also indicated changes in locomotion, which were included as a sign of pain and suffering. The QoL assessment by the expert was a reliable source of information. However, the reliability of QoL assessments can be dependent on the circumstances, so improving the efficacy of both qualitative and quantitative behavioral analyses is important for future studies.

In conclusion, our study documented the behavioral characteristics of an extremely old individual rhesus macaque and identified physical and cognitive decline. While physical decline may exacerbate cognitive difficulties, the two processes are distinct and require separate attention in both monitoring and treatment. We hope that future studies will consider the behavioral symptoms observed in this study as monitoring items to better understand physical and cognitive decline in primates. Compared with physical decline, reports on the relationship between cognitive decline and well-being of primates are still scarce. Future studies are warranted on inclusion of changes in cognitive needs while considering well-being of animals as well as their cognitive enrichment.

## Acknowledgement

This study was financially supported by Nakatsuji Foresight Foundation 2021, Murata Science Foundation (M23-032), TOYOTA foundation (D23-R-0028) and Japan Society for Promotion of Science (24K02112 and 24K00167). We thank Noah T. Dunham for the kind advice on gait analyses. We also thank Hiroki Ishikawa, Michiko Fujisawa, Nobuaki Yoshida and the staff of the Kyoto City Zoo for their help in this study. ChatGPT-4o was used to check the English of this paper and to assist in generating R codes for creating figures.

## Statement of Contribution

YY and HB conceived the study. YY and DN collected the data. YY analyzed the data. YY drafted the manuscript. All authors reviewed and approved the final manuscript.

## Data availability

The data that support the findings of this study are openly available in Open Science Framework (Yamanashi, 2024).

## Conflict of interest disclosure

The authors declare no conflicts of interest.

## Biosketch

Yumi Yamanashi is the principal researcher at Kyoto City Zoo, where she mainly works in the area of animal welfare. Haruna Bando and Keita Niimi are keepers, and Daisuke Nakagawa is a veterinarian at Kyoto City Zoo. Susumu Iwaide and Tomoaki Murakami are researchers in animal pathology, with expertise in amyloidosis and related diseases.

---

**Supplementary table 1.** The subjects of the study.

| ID | Gender | Date of birth | Age* | Reference date |
| --- | --- | --- | --- | --- |
| Isoko | Female | 1978/4/15 | 43* | 2021/9/12 |
| Gongo | Female | 1990/5/5 | 32* | 2022/9/30 |
| Beat | Female | 1985/4/1 | 37* | 2022/12/26 |
| Tono | Male | 1997/5/12 | 26 | 2023/5/30 |
| Cheeko | Female | 1997/5/22 | 26 | 2023/5/30 |
| Tsubone | Female | 1998/6/7 | 24 | 2023/5/30 |
| Tenpura | Female | 1999/5/11 | 24 | 2023/5/30 |
| Boko2 | Female | 1999/8/6 | 23 | 2023/5/30 |
| Kankan | Female | 2001/5/16 | 22 | 2023/5/30 |
| Nano | Female | 2002/5/15 | 21 | 2023/5/30 |
| Saeko | Female | 2003/6/7 | 19 | 2023/5/30 |
| Agi | Female | 2003/6/15 | 19 | 2023/5/30 |
| Koyuriko | Female | 2004/8/25 | 18 | 2023/5/30 |
| Yoko | Female | 2006/4/29 | 17 | 2023/5/30 |
| Yone | Female | 2008/8/28 | 14 | 2023/5/30 |

\*The ages of the individuals were calculated using the data from the reference date column. The ages of three individuals (Isoko, Gongo, and Beat) were determined based on their dates of death, while the ages of the others were calculated at the time of the study.

**Supplementary table 2.**
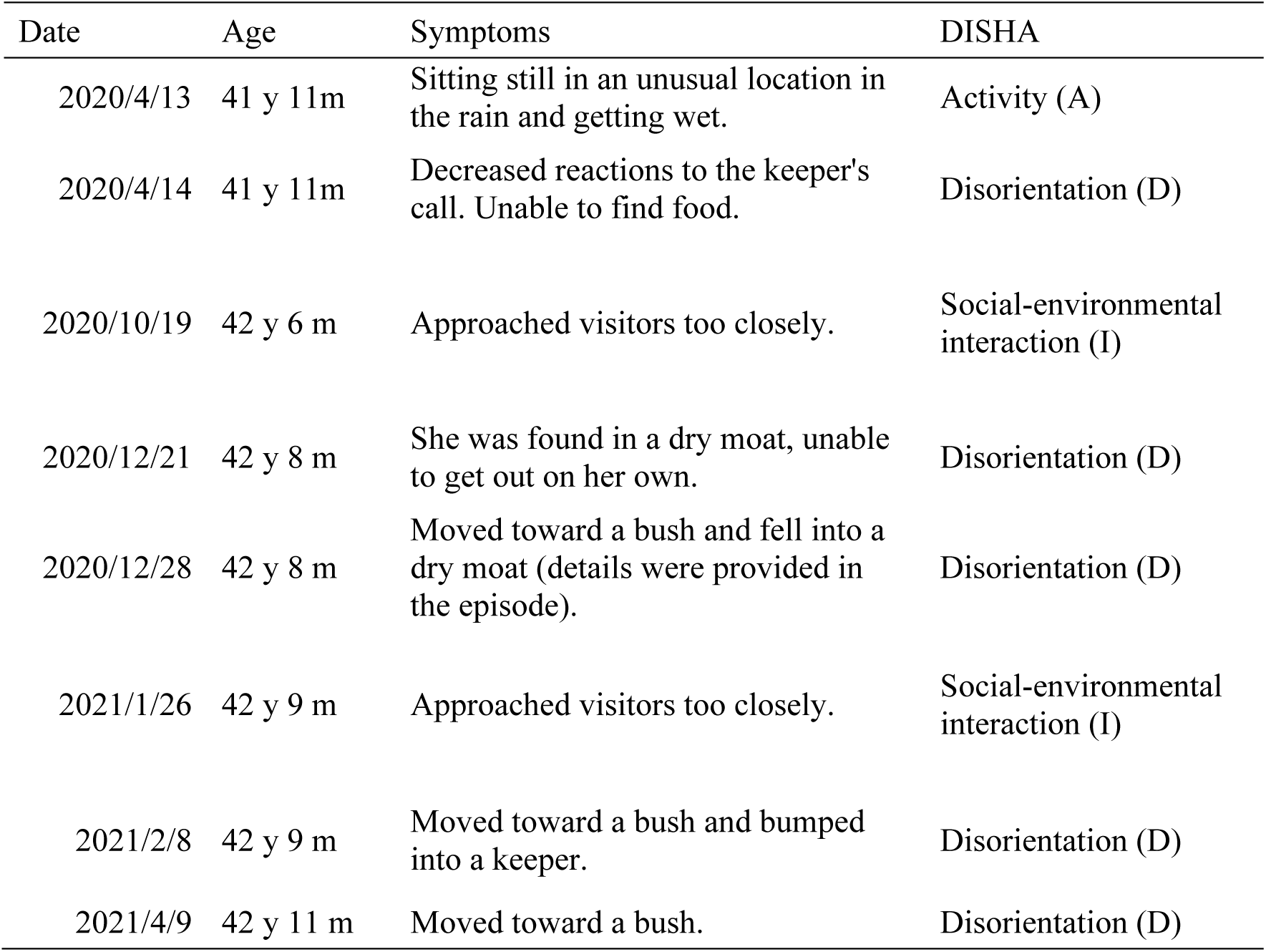
The extracts from the keepers’ diary and their possible relationships with DISHA.

**Supplementary table 3.**
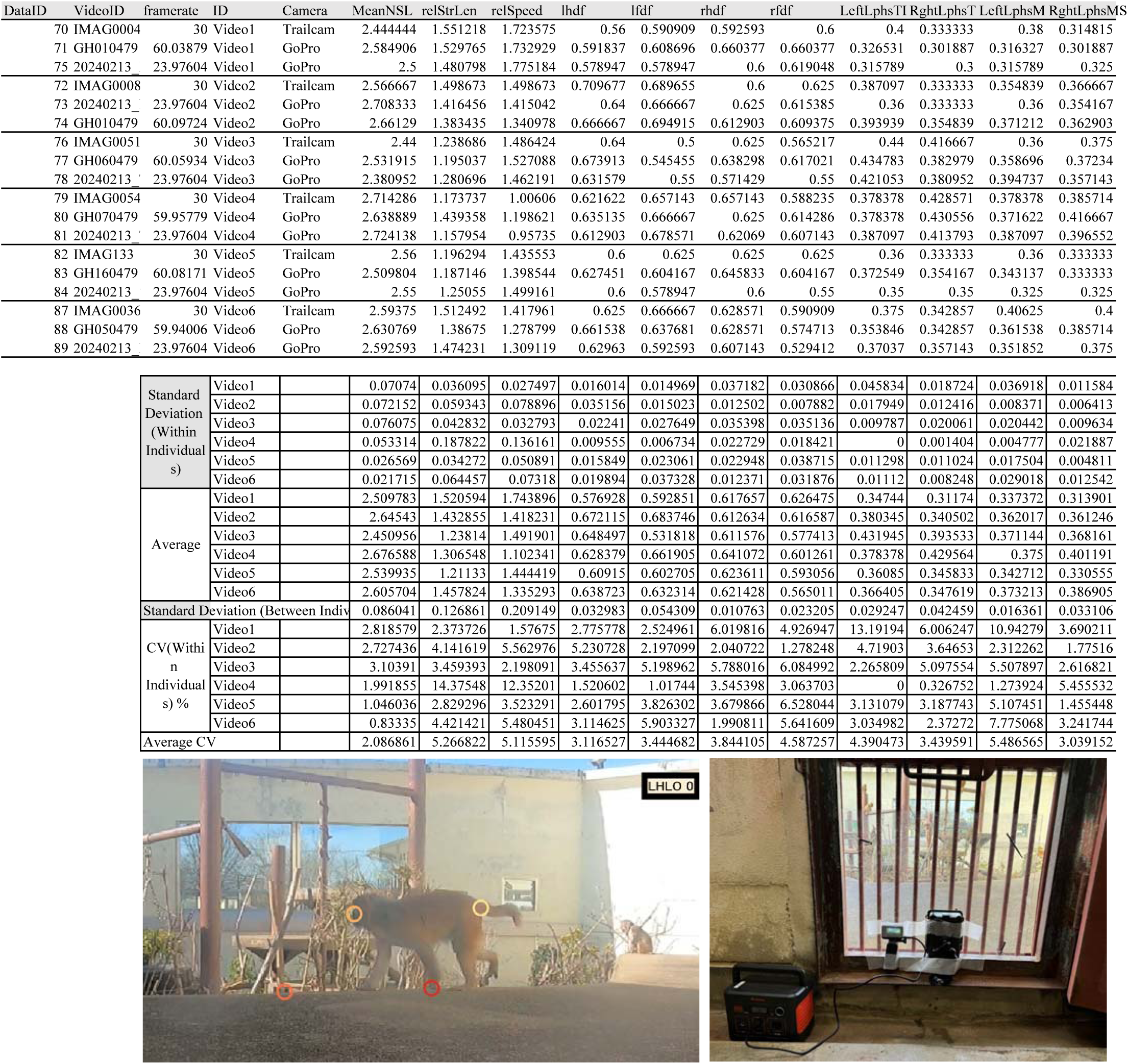
The results of gait analyses obtained from different cameras.

**Supplementary table 4.** The Quality of Life (QoL) assessment sheet used in the study.

| Category | 5 | 4 | 3 | 2 | 1 |
| --- | --- | --- | --- | --- | --- |
| Pain | No difficulty noted getting up and down | Slight difficulty getting up and down | Moderate difficulty getting up. May take 2-3 attempts to get up | Significant difficulty getting up. Makes more than 3 attempts to get up or spends most of the time on its knees. Appears uncomfortable | Extreme difficulty getting up. Unable to and animal stops trying to after several attempts |
| Daily Maintenance | Eating, drinking, grooming, urinating and defaecating normally. Good body condition | Normal maintenance behavior. Body condition fair | Normal maintenance behavior. Poor body condition | Not interested in grooming but still eating, drinking, urinating and defaecating normally. Poor body condition | Not interested in grooming, eating and drinking. Poor body condition |
| Appetite | Interested in food or eats 100% of food | Consumed immediately or eats 75% of food | Consumed after several hours or eats 50% of food. Special food preparation required | Investigated but did not consume or eats 25% of food. Special food preparation required | Not interested in food or not eating at all |
| Movement | No noticeable lameness. Fluid movement | Slight lameness, intermittent. Taking shorter steps | Moderate lameness, noticeable all the time but animal is bearing weight. Noticeably stiff most of the time | Significant lameness, noticeable all the time and animal holds limb up when stationary or when moving. Shuffling and animal appears uncomfortable | Severe lameness, non-bearing lameness up all the time. Animal not willing to move around |
| Interaction with conspecifics/ keepers / environmental stimulus | High interest to interact (with others or the environment). Social status maintained (for social animals) | Moderate to high interest to interact. Social status maintained | Moderately interested to interact. Social status maintained | Very little interest shown to interact. Social status not maintained | No signs of interest to interact. Social status not maintained |
| Manipulation | No noticeable concerns. The animal can grasp objects and explore normally. | Some mild clinical signs, such as slight tremors when grasping objects but there are no issues with its ability to grasp or explore. | Tremors when grasping objects, and there are some difficulties in grasping and exploring. The animal occasionally fails. | Tremors when grasping objects, and there are difficulties in grasping and exploring. The animal fails about 80% of the time, making it challenging to perform these actions independently. | Completely unable to grasp objects or explore on its own. |
| Veterinary findings | No diseases, injuries, or disabilities at all. | Diseases, injuries, or disabilities are present, but they are curable | Incurable diseases, injuries, or disabilities are present, but they do not interfere with daily life. | Incurable diseases, injuries, or disabilities are present, causing some difficulties in daily life. | Incurable diseases, injuries, or disabilities are present, causing significant difficulties in daily life. |

## Notes

https://osf.io/4kjsz/overview?view_only=eb67279f3820486eb8e434cf29de1795

### Competing Interest Statement

The authors have declared no competing interest.

